# Anti-Asialo GM1 treatment during secondary *Toxoplasma gondii* infection is lethal and depletes T cells

**DOI:** 10.1101/550608

**Authors:** Daria L. Ivanova, Steve L. Denton, Jason P. Gigley

## Abstract

Using vaccine challenge model of *T. gondii* infection, we found that treatments with two commonly used for NK cell depletion antibodies resulted in different survival outcomes during secondary infection. Anti-ASGM1 resulted in 100% death and greater parasite burden at the site of infection than anti-NK1.1. Anti-NK1.1 treatment resulted in increased parasite burdens, but animals did not die. Further we found that anti-ASGM1 treatment depleted T cells. CD8+ T cells were more susceptible that CD4+ T cells to the treatment. ASGM1 was expressed on a higher percentage of CD8+ T cells than CD4+ T cells and CD8+ T cells. In *T. gondii*-immunized animals ASGM1 was enriched on effector memory (Tem) and central memory (Tcm) CD8+ T cells. However, Tem were more susceptible to the treatment. After secondary infection, Tem, Tcm, effector (Tef) and naïve (Tn) CD8+ T cells were positive for ASGM1. Anti-ASGM1 treatment during reinfection resulted in greater depletion of activated IFNγ+, Granzyme B+, Tem and Tef than Tcm and Tn CD8+ T cells. Anti-ASGM1 also depleted IFNγ+ CD4+ T cells. Recombinant IFNγ supplementation prolonged survival of anti-ASGM1 treated mice, demonstrating that this antibody eliminated IFNγ-producing T and NK cells important for control of the parasite. These results highlight that anti-ASGM1 antibody is not an optimal choice for targeting only NK cells and more precise approaches should be used. This study uncovers ASGM1 as a marker of activated effector T cells and the potential importance of changes in sialylation in lipid rafts for T cell activation during *T. gondii* infection.

## Introduction

*T. gondii* is an obligate intracellular protozoan that infects ∼25-30% of people worldwide (Flegr et al., 2014, Jones and Dubey, 2012). Infection with the parasite in people is one of the leading causes of foodborne illnesses in the U.S. resulting in hospitalization (Scallan et al., 2011). Despite induction of a robust CD8+ T cell response, the parasite does not get completely cleared and establishes a persistent infection in the brain and muscles of the host (Harker et al., 2015, Wohlfert et al., 2017). Recent studies suggest that infection with the parasite correlates with schizophrenia, depression, behavioral changes, and neurodegenerative disorders (Lang et al., 2018, Donley et al., 2016). In the patients with weakened immune system, the *T. gondii* parasites can cause toxoplasmic encephalitis and death (Luft and Remington, 1992, Kodym et al., 2015). Currently, there are no vaccines or treatments that completely clear the infection (Radke et al., 2018). Understanding how the immune system functions in response to the parasite is paramount for the development of novel therapeutic approaches to treat this infection.

*T. gondii* infection induces a robust Th1 response that provides long-term protection through the production of IFNγ by CD8+ T cells (Suzuki et al., 1988, Gazzinelli et al., 1991). However, IFNγ is also produced early during infection by innate immune cells, including NK cells, ILC1 and neutrophils (Sher et al., 1993, Klose et al., 2014, Sturge et al., 2013). The approaches taken to define the role of different innate immune cell populations have included cell depletion studies *in vivo* using antibodies to the different cell populations and animals with mutations or genetic deficiencies that result in the absence of the cell of interest. REF These approaches of course are not perfect and have off target effects making it difficult to fully understand the role of different cell types in a disease. This situation has become more apparent for studies investigating NK cells and depends upon the disease model being studied (Nishikado et al., 2011). NK cells are usually depleted using NK cell specific antibodies *in vivo* (Victorino et al., 2015). However, in some disease models, antibody depletion of NK cells either targeted NK cells alone, or targeted additional subsets of immune cells (Nishikado et al., 2011). The conventional approach to target NK cells *in vivo* utilizes antibodies against asialo GM1 (ASGM1) or the surface marker NK1.1 (Young et al., 1980). Anti-ASGM1 antibody reacts with a glycosphingolipid expressed on the surface NK cells, basophils, and on some subsets of γδT, NKT, CD8 T cells and macrophages (Nishikado et al., 2011, Slifka et al., 2000, Wiltrout et al., 1985, Kasai et al., 1980). In contrast, anti-NK1.1 targets a glycoprotein expressed on NK cells and some subsets of γδT, NKT, and CD8+ T cells (Slifka et al., 2000). Anti-NK1.1 depletion of NK cells is limited in specific mouse strains due to an allelic variation in the NKR-P1 gene that encodes NK1.1. REF Therefore, anti-NK1.1 can only be used in C57BL/6, SJL, and NZB and not in other strains (BALBC etc.) (Mesci et al., 2006, Carlyle et al., 2006). Anti-ASGM1 is an alternative and is very commonly used for NK cell depletion because it effectively depletes NK cells *in vivo* in many mouse strains and other species. REF Although there are mouse strains with genetic alterations that result in NK cell deficiencies without the use of antibodies, they have additional defects that make interpretation of studies difficult (Jessen et al., 2011, Eckelhart et al., 2011). Therefore, when choosing between the use of antibodies for NK cell depletions and genetically modified NK cell deficient mice, many investigators lean towards the use of the antibodies due to their availability, ease of use, and the ability to have a temporal control of cell ablation *in vivo*.

Many studies have demonstrated that NK cells are essential for protection against *T. gondii* infection by using anti-ASGM1 or anti-NK1.1 treatments (Denkers et al., 1993, Gazzinelli et al., 1993, Goldszmid et al., 2012, Askenase et al., 2015). While there is no doubt about NK cell importance, whether these experimental approaches had additional effects on the immune response to the parasite is not clear. In this study we found that anti-NK1.1 and anti-ASGM1 significantly differed in their impact on mouse survival and parasite burdens when administered during lethal *T. gondii* challenge of mice vaccinated with the attenuated type I RH strain *cps1-1* (CPS {Fox, 2002 #390}). The parasite burdens were higher and the survival rate was reduced after the treatment with anti-ASGM1 as compared to anti-NK1.1. We further explored the reason for this difference and discovered that while both antibodies depleted NK cells, anti-ASGM1 also significantly depleted CD8+ and CD4+ T cells both in spleen and the site of challenge infection. Anti-NK1.1 treatment did not have as dramatic of an effect. Interestingly, CD8+ T cells were more affected by anti-ASGM1 treatment than CD4+ T cells. ASGM1 was expressed on the surface of CD4+ and CD8+ T cells in CPS-vaccinated animals. Further dissection of whether ASGM1 was expressed more on memory T cells after vaccination revealed that effector memory (Tem, CD62L-CD44+) and central memory (Tcm, CD62L+CD44+) CD8+ T cells both had higher concentrations of ASGM1 on their surface. However, during secondary challenge of vaccinated mice, all T cell subsets, including memory (Tem and Tcm), effector (Tef) and naïve (Tn) cells increased ASGM1 on their surface. Interestingly, anti-ASGM1 treatment during RH challenge of CPS-vaccinated animals effectively eliminated most Tem and Tef (CD62L-) CD8+ T cells leading to an increase in naïve CD8+ T cells. Anti-ASGM1 significantly reduced the frequency and number of activated IFNγ+, Granzyme B+ and polyfunctional IFNγ+GranzymeB+ CD8+ T cells and IFNγ+ CD4+ T cells in RH challenged animals. Recombinant IFNγ supplementation of depleted animals prolonged their survival. These studies demonstrate that NK cell depletion with anti-ASGM1 during *T. gondii* infection also eliminates CD8+ and CD4+ T cells in *T. gondii* vaccinated animals and that ASGM1 may be important in the activation of T cells important in control of parasite infection.

## Results

### Anti-NK1.1 and anti-ASGM1 have different effects on control of T. gondii infection

The importance of NK cells for early control of *T. gondii via* IFNγ is well established (Denkers et al., 1993, Sher et al., 1993, Combe et al., 2005, Goldszmid et al., 2012). We recently demonstrated that NK cells are also important in controlling secondary parasite infection in a vaccine-challenge model of *T. gondii* (Ivanova D. et al. in prep.). Our experimental approach to testing the importance of NK cells in this model involved the use of NK cell specific antibodies anti-NK1.1 and anti-ASGM1 to deplete the cells *in vivo*. C57BL6 mice were immunized with the attenuated type I strain *cps1-1* (CPS) and five weeks later challenged with a lethal dose of the CPS parental parasite strain RH and mouse survival and parasite burdens by real time semi quantitative PCR measured. We observed that anti-ASGM1 treatment resulted in the death of all of the animals within 11 days of type I RH challenge and 31 days after type II ME49 challenge (Fig. 1A). However, Anti-NK1.1 treatment did not result in death after type I RH and type II ME49 challenge (Fig. 1A). We also observed that parasite burdens were higher in immunized then challenged mice after anti-ASGM1 treatment (Fig. 1B-C, Supplementary Fig. 1A-C). In addition, the parasite burdens were greater in the mice treated with anti-ASGM1 compared to anti-NK1.1 (Supplementary Fig. 1B-C). In a separate set of experiments mice were chronically infected with ME49 cysts and then challenged with RH tach. in the presence or absence of anti-ASGM1. Anti-ASGM1-treated animals died much sooner compared to untreated controls (Fig. 1D). These results showed that, in comparison with anti-NK1.1 treatment, the administration of anti-ASGM1 during secondary *T. gondii* infection had a more robust effect on the parasite dissemination and the death of mice. Since both antibody treatment regimens effectively depleted NK cells (Supplementary Fig. 1D), this result suggested that anti-ASGM1 may have depleted additional cell subsets necessary for the protection during secondary *T. gondii* infection.

**Figure 1.**
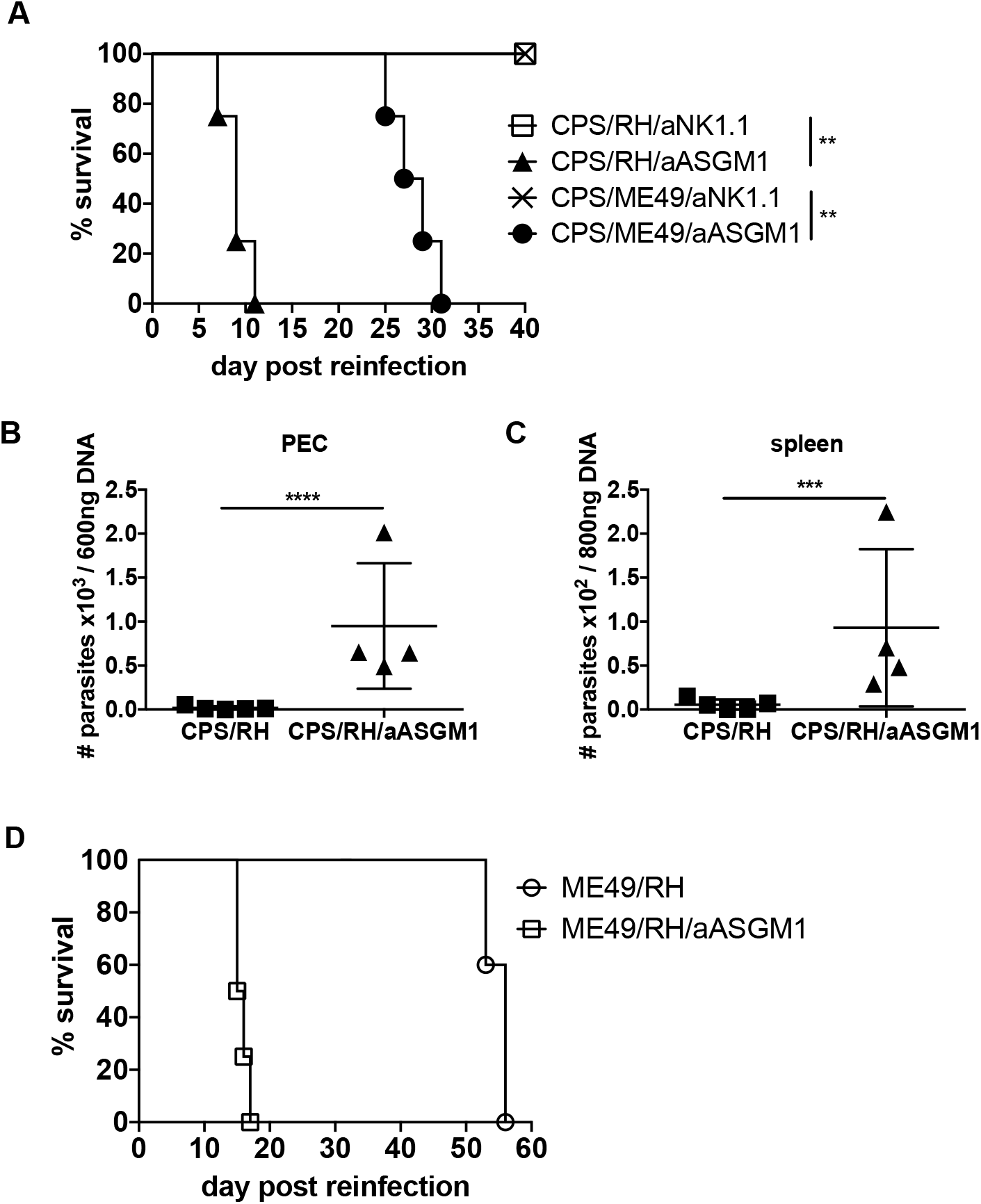
Anti-ASGM1 treatment during secondary *T. gondii* infection leads to rapid death and increased parasite burdens. (**A-C**) Immunized i.p. with 1 × 10^6^ CPS tach. C57BL/6 mice were five to six weeks later infected i.p. with 1 X10^3^ RH tach. (**A-C**) or i.g. with 100 ME49 brain cysts (**A**) and treated with anti-NK1.1 (PK136) (**A**) or anti-ASGM1 (**A-C**) during secondary infections. (**A**) The survival curve represents one of three to four experiments, n=4 to 5. (**B** and **C**) The parasite burdens measured by real time PCR for *T. gondii* B1 gene 8 days after RH infection in PEC (**B**) and spleen (**C**). The data represent one of two experiments, n=4 to 5. (**D**) Infected with 10 ME49 cysts i.p. C57BL/6 mice were four weeks later infected with 1 × 10^3^ RH tach. i.p. and treated with anti-ASGM1 upon reinfection. The survival curve represents one of two experiments, n=3 to 5. The log-rank (Mantel-Cox) test was used to evaluate survival rates. Ordinary one-way ANOVA was used to evaluate the parasite burdens. Data are mean ± SD. *p<0.05, **p<0.01, ***p<0.001, ****p<0.0001.

### Anti-ASGM1 treatment depletes CD8+ and CD4+ T cells after secondary challenge of T. gondii immunized mice

*In vivo* antibody depletion of cells can have unexpected off target effects on other immune cells (Nishikado et al., 2011). Several studies showed that T cell numbers were reduced after anti-ASGM1 treatment during certain viral infections, including vaccinia virus, LCMV, reovirus and RSV (Stitz et al., 1986, Doherty and Allan, 1987, Parker et al., 1988, Moore et al., 2008). One study showed T cell numbers were reduced and the parasite burdens were increased after anti-ASGM1 treatment, however, no death was observed and it was not clear whether the parasite burdens were increased due to the depletion of NK, T or both cell types (Moore et al., 2008). Based on these previous findings, increased susceptibility of *T. gondii* immunized and anti-ASGM1 treated during reinfection mice could be associated with the depletion of T cells. To address whether anti-ASGM1 depleted not only NK cells, but also T cells in CPS-immunized animals, the CD8+ and CD4+ T cell frequencies and absolute numbers were measured by flow cytometry in the peritoneum (PEC) and spleens of antibody treated or not treated immunized mice. Anti-ASGM1 treatment did not significantly affect T cells in the peritoneum, but the numbers of CD4+ and CD8+ T cells were significantly reduced in the spleen (Fig. 2A-E), demonstrating that in the absence of secondary challenge anti-ASGM1 also depleted T cells. To address whether this treatment also depleted CD4+ and CD8+ T cells during secondary challenge, anti-ASGM1 treated or non-treated CPS-immunized mice were challenged with RH and T cell numbers measured. Anti-ASGM1 administered upon RH challenge significantly reduced CD8+ T cell frequency and numbers in both peritoneum and the spleen compared to non-treated animals (Fig. 2A-E). In contrast, CD4+ T cells were not significantly affected by anti-ASGM1 treatment during challenge infection. In contrast to anti-ASGM1, anti-NK1.1 treatment did not reduce CD8+ and CD4+ T cell frequency and numbers in both peritoneum and spleen (Supplementary Fig. 2). This result suggests that the increased susceptibility of mice during anti-ASGM1 treatment upon secondary *T. gondii* infection could be associated with the depletion of CD8+ and CD4+ T cells.

**Figure 2.**
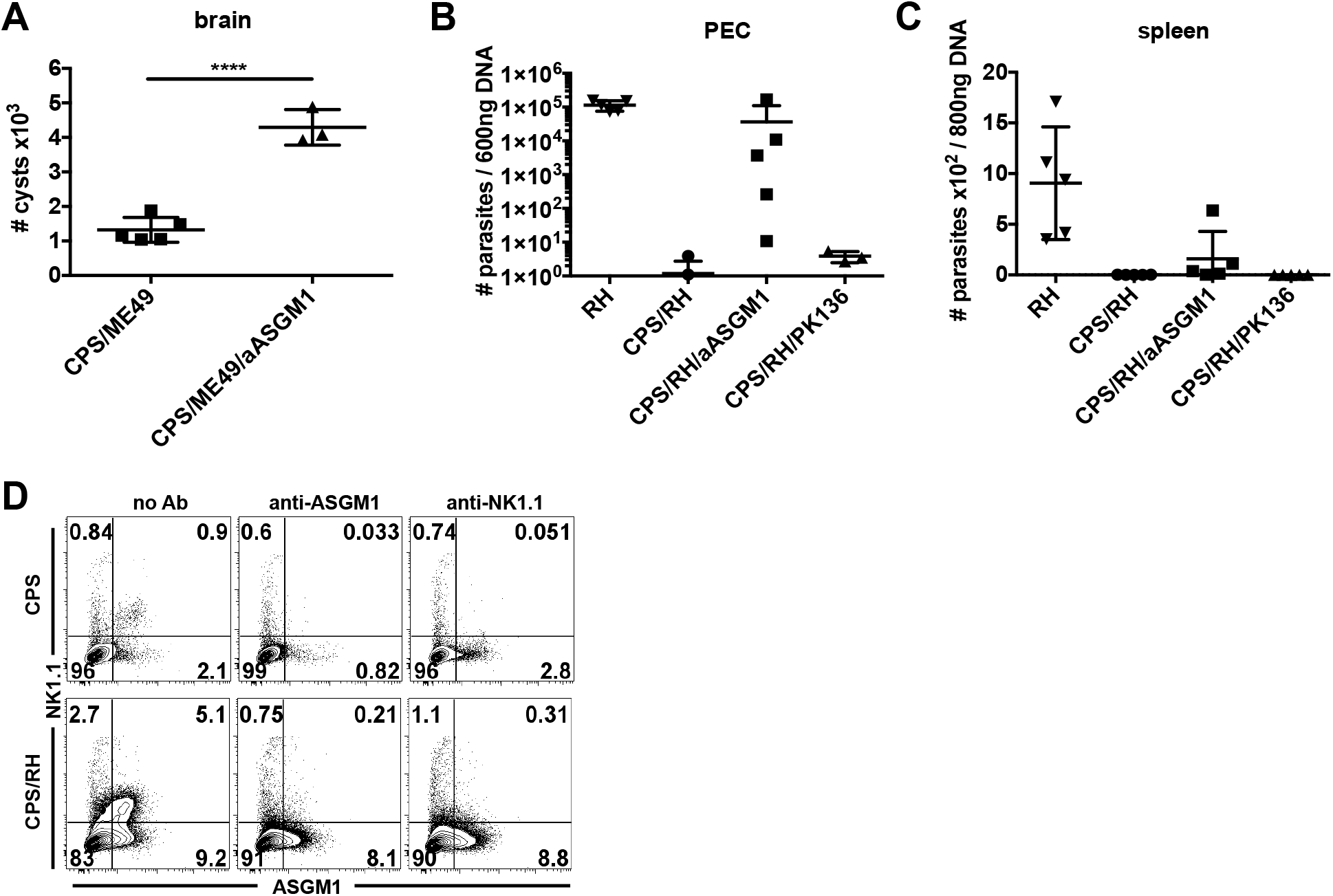
Anti-ASGM1 and anti-NK1.1 treatments during secondary *T. gondii* infection. (**A-D**) Immunized i.p. with 1 × 10^6^ CPS tach. C57BL/6 mice were five to six weeks later infected i.g. with 20 ME49 brain cysts (**A**) or i.p. with 1 × 10^3^ RH tach. (**B-D**) and treated with anti-ASGM1 (**A-D**) or anti-NK1.1 (PK136) (**B-D**) during secondary infection. (**A**) The brain cysts were counted after ME49 reinfection at 35 dpi in control mice and immediately after the death in anti-ASGM1 treated mice. The data represent 1 experiment, n=3 to 5. (**B-C**) The parasite burdens were measured by real time PCR for *T. gondii* B1 gene 8 days after RH reinfection in PEC (**B**) and spleen (**C**). The data represent one experiment, n=3 to 5. (**D**) NK cell (gated as NK1.1+ASGM1+ live lymphocytes) depletion after anti-ASGM1 or anti-NK1.1 treatment was measured by flow cytometry. Representative contour plots from one of two experiments, n=3 to 4. Ordinary one-way ANOVA, data are mean ± SD, ****p<0.0001.

### ASGM1 is differentially expressed on CD8+ and CD4+ T cells in T. gondii immunized mice

The previous result showed that anti-ASGM1 treatment depleted CD8+ and CD4+ T cells with potentially CD8+ T cells being targeted more than CD4+ T cells. Previous studies indicate that CD8+ and CD4+ T cells have higher levels of ASGM1 on their surface after viral infections (Slifka et al., 2000, Moore et al., 2008) possibly making them susceptible to anti-ASGM1 depletion. Therefore, we measured by flow cytometry whether ASGM1 was expressed on CD8+ and CD4+ T cells after *T. gondii*-immunization in mice and how ASGM1 levels changed on T cells after challenge infection. ASGM1 was detected on both cell types in immunized mice (Fig. 3A-F). After secondary parasite infection, the frequency and number of ASGM1+ CD4+ T cells and number of ASGM1+ CD8+ T cells increased at the challenge site (Fig. 3C-E). The frequency of ASGM1+ CD8+ T cells in peritoneum and spleen did not increase (Fig. 3 A-D). Interestingly, ASGM1 was expressed on a much higher percentage of CD8+ T cells than CD4+ T cells in both PEC and spleen.

**Figure 3.**
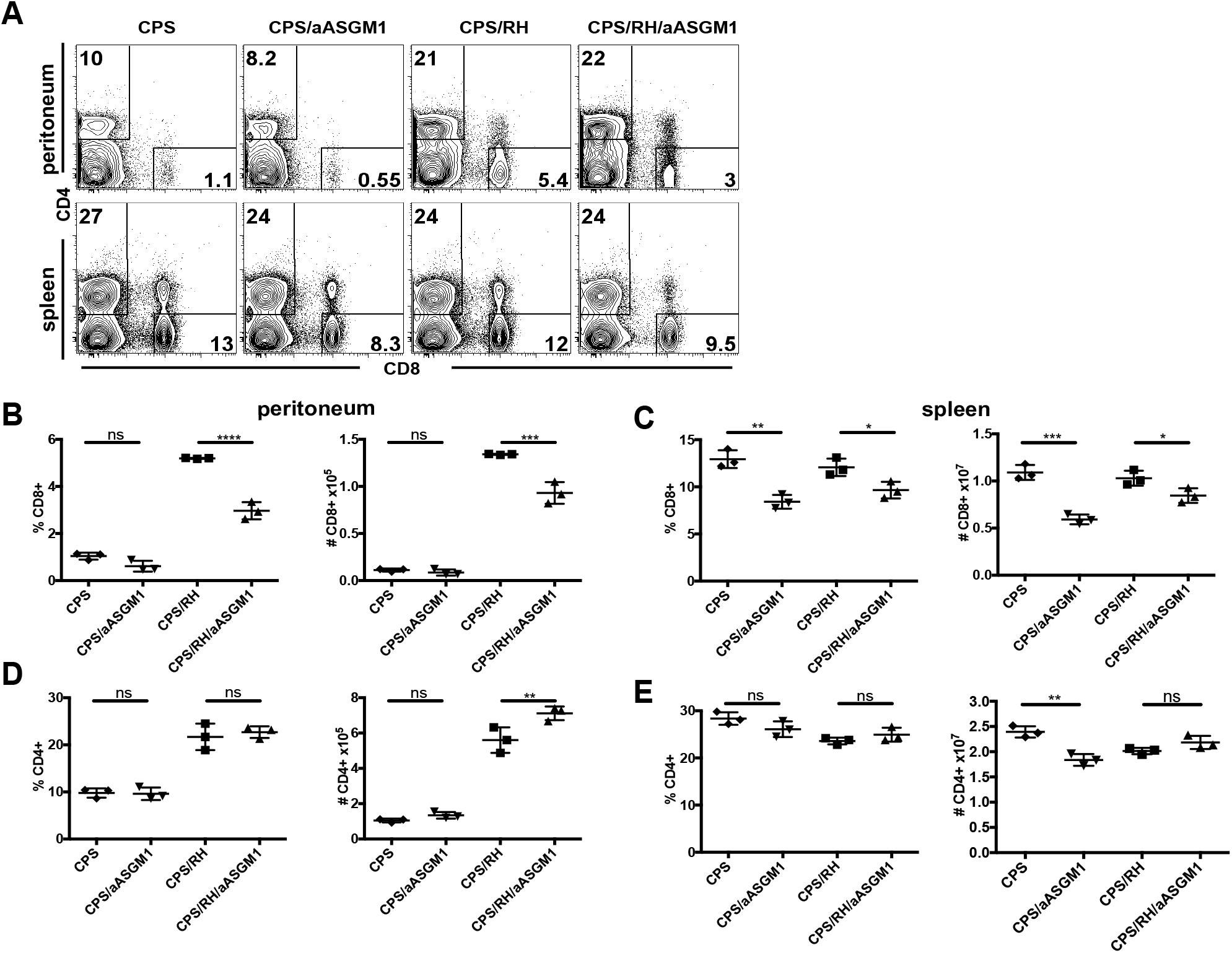
Anti-ASGM1 treatment depletes CD8+ and CD4+ T cells in *T. gondii*-immunized mice. **(A-E)** C57BL/6 mice were immunized i.p. with 1 × 10^6^ CPS tach. and infected i.p. with 1 × 10^3^ RH tach. and treated with anti-ASGM1 antibody. PECs and spleens were analyzed by flow cytometry at day 3 post RH infection. (**A**) Representative contour plots of CD8+ and CD4+ T cells (gated on NK1.1-live lymphocytes). The percentages and numbers of CD8+ T cells per PEC (**B**) or spleen (**C**). The percentages and numbers of CD4+ T cells per PEC (**D**) or spleen (**E**). Data are representative of one of three experiments, n=3 to 4. Data are mean ± SD. *p<0.05, **p<0.01, ***p<0.001, ****p<0.0001, ordinary one-way ANOVA.

### Anti-ASGM1 treatment depletes effector and effector memory CD8+ T cell subsets

Previous studies reported that ASGM1 was differentially enriched among different CD8+ T cell subsets. In naïve unimmunized mice central memory phenotype (CD44^high^, CD62L+, CCR7+ and CD122+) CD8+ T cells expressed more ASGM1 + than other T cell subsets (Kosaka et al., 2006, Kosaka et al., 2007). Since the mice in our studies were immunized and the protection was lost when the animals were treated with anti-ASGM1, this could suggest that ASGM1 might be enriched on *T. gondii* specific memory T cell populations. Therefore, we next determined whether ASGM1 was expressed on certain subsets of CD8+ T cells and whether these same subsets were preferentially depleted by anti-ASGM1 treatment. CPS-immunized mice were treated with anti-ASGM1 in the absence or presence of RH infection and T cell phenotype was assayed based on the expression of CD44, a marker of antigen-experienced T cells (Budd et al., 1987) and CD62L, a marker of cells that home to lymph nodes (Lefrancois, 2006). In CPS-immunized mice CD8+ T cells were distributed between Tcm (central memory, CD44+CD62L+) > Tn (naïve, CD44-CD62L+) > Tem (effector memory, CD44+CD62L-) > Tef (effector, CD44-CD62L-) (Fig. 4A) (Srivastava et al., 2018). ASGM1 was mainly expressed on antigen-experienced CD44^high^ Tcm and Tem CD8+ T cells in CPS immunized mice (Fig. 4B). After infection of CPS-immunized mice with RH, ASGM1 was also expressed on Tef and Tn CD8+ T cells (Fig. 4B). Anti-ASGM1 reduced the frequency of all of the CD8+ T cell subsets in CPS-immunized animals without RH challenge, except for the Tn subset, which increased after the depletion. After RH challenge, there was an expected increase in Tef. However, anti-ASGM1 treatment during this challenge infection significantly reduced effector T cell populations, including Tef and Tem, while the frequency of naïve T cells increased. Interestingly, among all T cell subsets, Tcm cells seemed resistant to anti-ASGM1 treatment. Our results indicate that ASGM1 is enriched on different CD8+ T cell subsets after CPS immunization and secondary infection. We also observe that anti-ASGM1 treatment preferentially depletes effector populations of CD8+ T cells including Tem and Tef.

**Figure 4.**
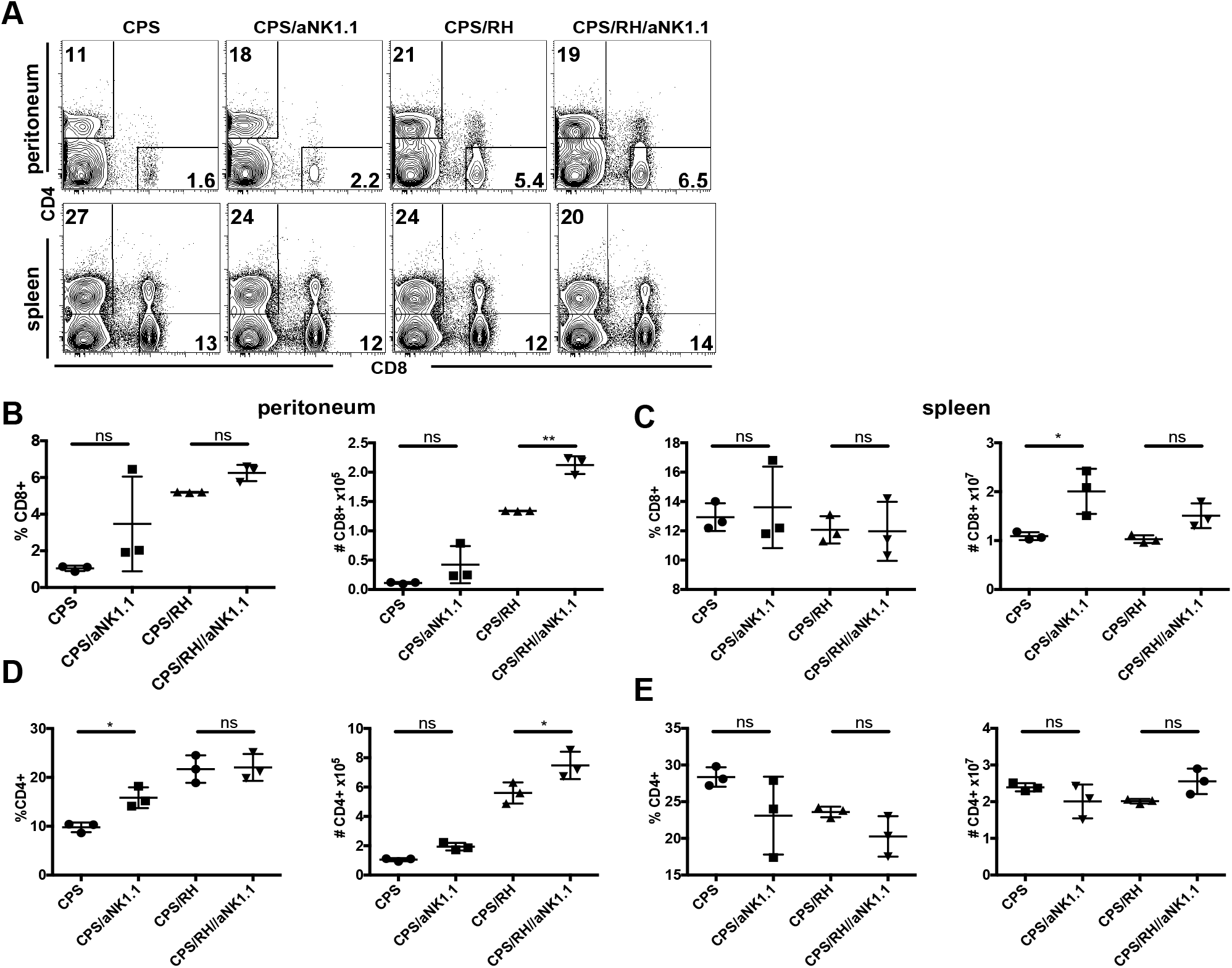
Anti-NK1.1 treatment does not deplete CD8+ and CD4+ T cells in *T. gondii*-immunized mice. C57BL/6 mice were immunized i.p. with 1 × 10^6^ CPS tach. and infected i.p. with 1 × 10^3^ RH tach. six weeks later and treated with anti-NK1.1 antibody. PECs and spleens were analyzed by flow cytometry at day 3 post RH infection. (**A**) Representative contour plots of CD8+ and CD4+ T cells (gated on NK1.1-live lymphocytes). The percentages and numbers of CD8+ T cells per PEC (**B**) or spleen (**C**). The percentages and numbers of CD4+ T cells per PEC (**D**) or spleen (**E**). Data are representative of one of three experiments, n=3 to 4. Data are mean ± SD. *p<0.05, **p<0.01, ***p<0.001, ****p<0.0001, ordinary one-way ANOVA.

CD11a and CD49d on CD8+ T cells can be used as surrogate markers of *T. gondii*-specific CD8+ T cells (Hwang et al., 2016). We therefore further explored whether *T. gondii*-specific CD11a+CD49d+ CD8+ T cells were depleted by anti-ASGM1 treatment and expressed ASGM1. The majority of CD8+ T cells were CD11a+CD49d+ and anti-ASGM1 treatment resulted in a depletion of these cells in CPS-immunized mice (Fig. 4C). Anti-ASGM1 also depleted these cells during secondary RH infection. Similarly to the previously observed increase in naïve CD8+ T cells, non-*T. gondii* specific CD11a-CD49d-CD8+ T cells increased after the depletion. In accordance to the depletion effect of anti-ASGM1, CD11a+CD49d+ CD8+ T cells were highly enriched in ASGM1 expression in CPS-immunized mice and in the presence of RH challenge (Fig. 4D). The result suggested that ASGM1 is enriched on *T. gondii*-specific CD8+ T cells and these cells were depleted by anti-ASGM1 treatment.

### Anti-ASGM1 treatment depletes activated CD8+ and CD4+ T cells

The above data identified that ASGM1 was expressed on a majority of CD8+ T cells that expressed surrogate markers of *T. gondii* specificity (CD11a and CD49d) and effector CD8+ T cells were depleted the most by anti-ASGM1 treatment. One study showed that ASGM1+ CD8+ T cells produced more IFNγ than ASGM1-CD8+ T cells upon stimulation with IL-12 (+IL-2) *in vitro* (Kosaka et al., 2006). IL-12 is produced after *T. gondii* infection and is essential for protection *via* induction of IFNγ (Gazzinelli et al., 1993, Hunter et al., 1995b, Yap et al., 2000). To find whether the increased susceptibility to the secondary infection during anti-ASGM1 treatment could be associated with the depletion of IFNγ-producing T cells, CPS-immunized mice were challenged with RH and their CD8+ and CD4+ T cell functionality was assessed after treatment with ani-ASGM1 compared to non-treated controls. Peritoneal CD8+ T cells were stimulated with TLA (Fig. 5A-B) or PMA/ionomycin (Fig. 5C-D) and stained intracellularly for IFNγ and Granzyme B and single function as well as polyfuncitonal responses were measured by flow cytometry. Anti-ASGM1 treatment significantly reduced the frequency and numbers of IFNγ+, Granzyme B+ and polyfunctional IFNγ+GranzymeB+ CD8+ T cells in the peritoneum (Fig. 5A-D). In addition, there was a significant reduction in frequency and number of IFNγ+ CD4+ T cells after anti-ASGM1 treatment at the site of infection. Thus, anti-ASGM1 treatment depletes activated CD4+ and CD8+ T cells during secondary challenge of immunized mice.

**Figure 5.**
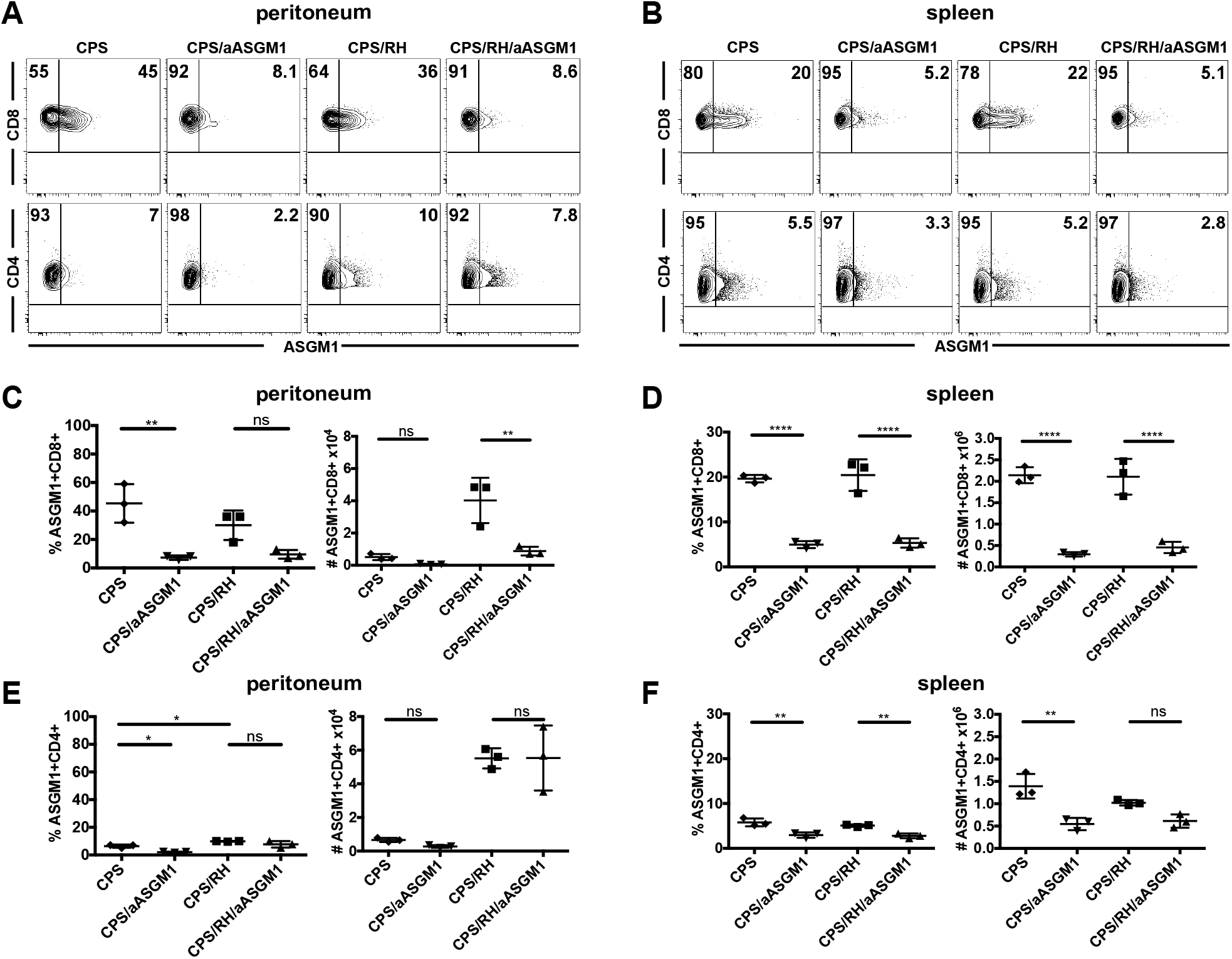
CD8+ and CD4+ T cells differentially express ASGM1 in *T. gondii* immunized mice. **(A-D)** C57BL/6 mice were immunized i.p. with 1 × 10^6^ CPS tach. and infected i.p. with 1 × 10^3^ RH tach. six weeks later. PECs and spleens were analyzed by flow cytometry at day 3 post RH infection. **(A)** Representative contour plots of ASGM1 expression on CD8+ and CD4+ T cells (gated on NK1.1-live lymphocytes) in peritoneum (**A**) or spleen (**B**). The percentages and numbers of CD8+ T cells per peritoneum (**C**) or spleen (**D**). The percentages and numbers of CD4+ T cells per peritoneum (**E**) or spleen (**F**). Data are representative of one of three experiments, n=3 to 4. Data are mean ± SD. *p<0.05, **p<0.01, ***p<0.001, ****p<0.0001, ordinary one-way ANOVA.

### IFNγ supplementation prolongs the survival of reinfected and anti-ASGM1 treated mice

Anti-ASGM1 treatment during reinfection increased the rate of mouse death and depleted IFNγ+ CD8+ and CD4+ T cells. To address whether the lethal effect of the antibody could be counterbalanced by IFNγ supplementation, recombinant IFNγ was administered during anti-ASGM1 treatment of CPS-immunized and RH infected mice. The cytokine supplementation had a therapeutic effect and significantly prolonged the survival of anti-ASGM1 treated mice (Fig. 6). Therefore, the detrimental effect of anti-ASGM1 treatment on the health and survival of mice could be associated with the depletion of IFNγ-producing CD8+ and CD4+ T cells in addition to IFNγ-producing NK cells.

**Figure 6.**
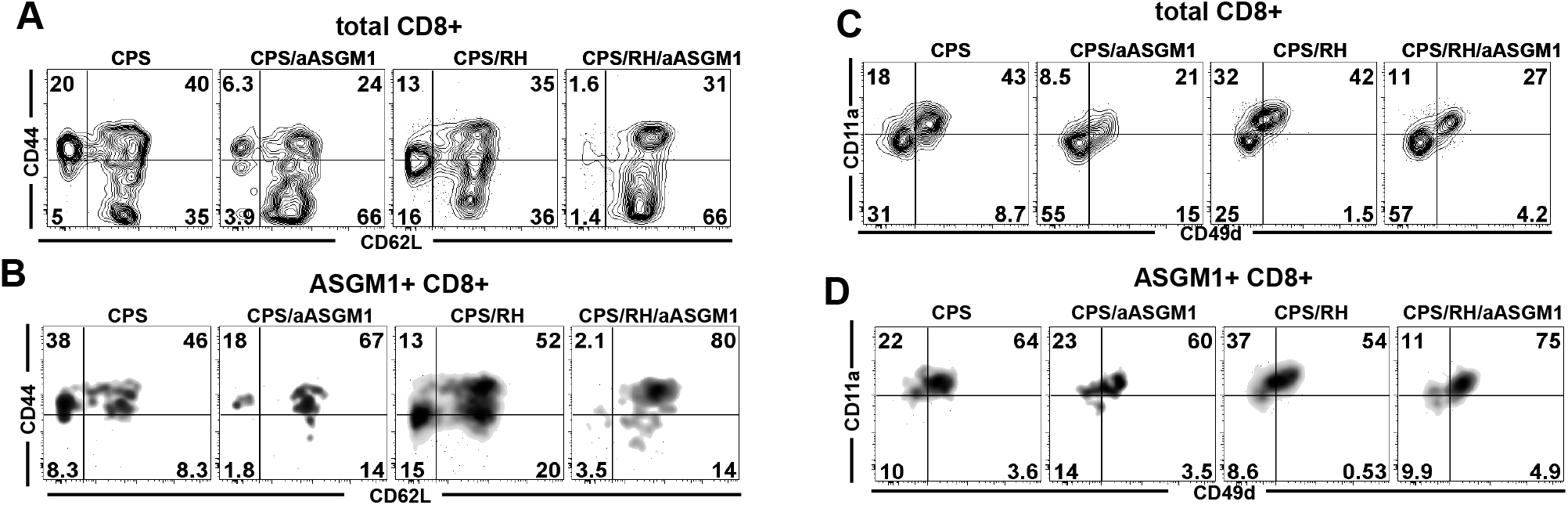
Anti-ASGM1 treatment depletes effector and effector memory CD8+ T cell subsets. (**A-D**) C57BL/6 mice were immunized i.p. with 1 × 10^6^ CPS tach. and infected i.p. with 1 × 10^3^ RH tach. six weeks later. PECs were analyzed by flow cytometry at day 3 post RH infection. Representative contour plots of CD44 and CD62L expression on total CD8+ T cells (gated on NK1.1-live lymphocytes) (**A**) and on ASGM1+ CD8+ T cells (**B**). Representative contour plots of CD11a and CD49d expression on total CD8+ T cells (gated on NK1.1-live lymphocytes) (**C**) and on ASGM1+ CD8b+ T cells (**D**). Data are representative of one of two experiments, n=3 to 4. Data are mean ± SD. *p<0.05, **p<0.01, ***p<0.001, ****p<0.0001, ordinary one-way ANOVA.

**Figure 7.**
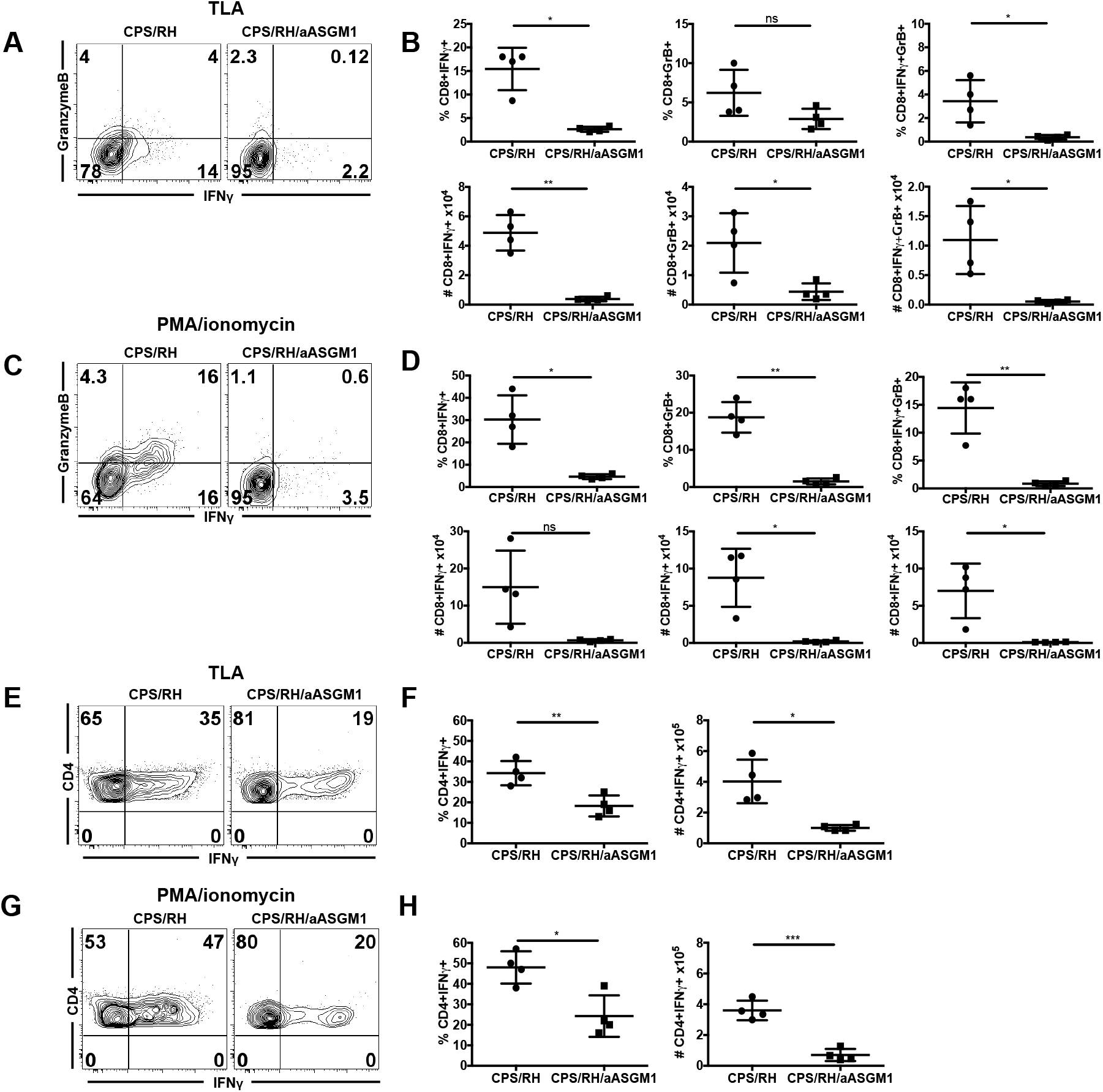
Anti-ASGM1 treatment depletes activated CD8+ and CD4+ T cells. (**A-H**) C57BL/6 mice were immunized i.p. with 1 × 10^6^ CPS tach. and infected i.p. with 1 × 10^3^ RH tach. six weeks later. PECs were analyzed by flow cytometry at day 3 post RH infection. Representative contour plots of intracellular staining for IFNγ and Granzyme B on CD8+ T cells (gated on CD8b+ live lymphocytes) after TLA stimulation (**A**) and in the presence of PMA/ionomycin stimulation (**C**). The percentages and numbers of IFNγ+, GranzymeB+ and IFNγ+GranzymeB+ CD8+ T after TLA (**B**) or PMA/ionomycin stimulation (**D**). Representative contour plots of intracellular staining for IFNγ on CD4+ T cells (gated on live lymphocytes) after TLA stimulation (**E**) and in the presence of PMA/ionomycin (**G**). The percentages and numbers of IFNγ+ CD4+ T after TLA (**F**) or PMA/ionomycin stimulation (**H**). Data are representative of 1 of 2 experiments, n=4 to 3. Data are mean ± SD. *p<0.05, **p<0.01, ***p<0.001, ****p<0.0001, ordinary one-way ANOVA.

**Figure 8.**
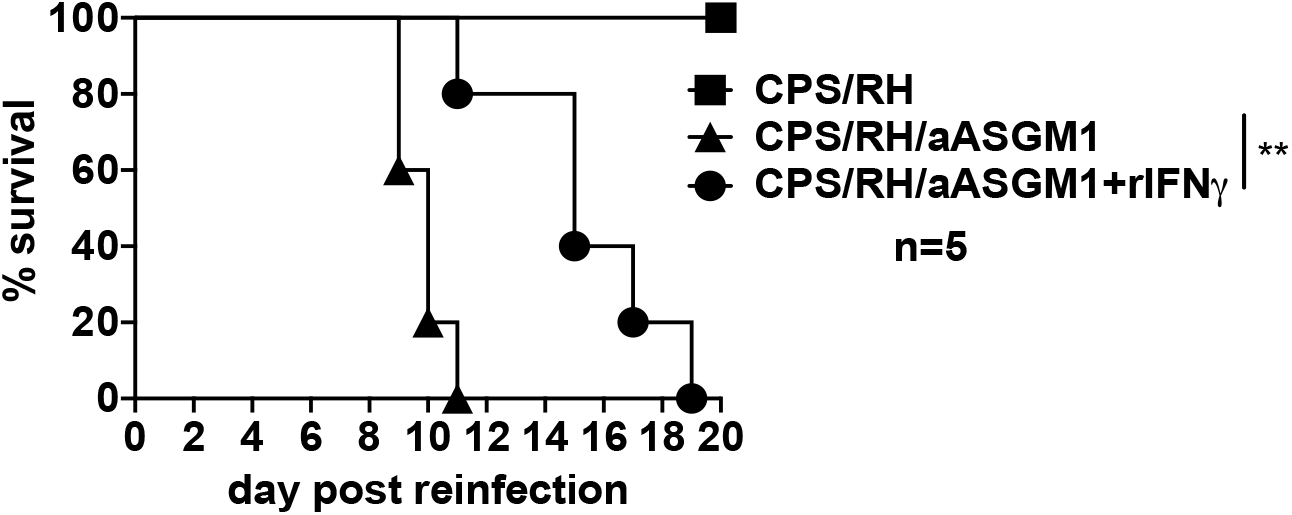
IFNγ supplementation prolongs the survival of *T. gondii* reinfected and anti-ASGM1 treated mice. Immunized i.p. with 1 × 10^6^ CPS tach. C57BL/6 mice were five to six weeks later infected i.p. with 1 × 10^3^ RH tach. and treated with anti-ASGM1 during secondary infection in the presence of recombinant IFNγ supplementation. The survival curve represents 1 experiments, n=5. The log-rank (Mantel-Cox) test was used to evaluate survival rates, **p<0.01.

## Discussion

Two commonly used for NK cell depletion antibodies exhibited different effects on the survival of mice infected with *T. gondii*. In contrast to anti-NK1.1, *in vivo* anti-ASGM1 treatment during secondary *T. gondii* infection led to a higher increase in parasite burdens and was lethal. This suggested that anti-ASGM1 could have an off-target effect, such as depletion of a cell type other than NK cells necessary for protection. In line with that, we found that anti-ASGM1 treatment resulted in the depletion of T cells. This is important because the anti-ASGM1 antibody is widely used to deplete NK cells (Ferreira et al., 2018, Haspeslagh et al., 2018). Many studies on the role of NK cells in *T. gondii* infection also relied on the use of anti-ASGM1 (Hunter et al., 1994, Haque et al., 1999, Goldszmid et al., 2012). Our data showed that this treatment is not a specific approach to address the role of NK cells during *T. gondii* infection. Our findings emphasize that anti-ASGM1 should be used with caution and in combination with other NK cell-specific methods.

ASGM1+ T cells could be essential for protection against *T. gondii* infection. A few previous studies showed that anti-ASGM1 treatment did not deplete T cells in naïve mice (Nishikado et al., 2011), but negatively affected T cells in the mice challenged with viruses (reovirus, lymphocytic choriomeningitis virus (LCMV), vaccinia, respiratory synovial virus (RSV)) and alloantigens (Stitz et al., 1986, Slifka et al., 2000, Moore et al., 2008). While the treatment was shown to deplete viral-activated T cells (Stitz et al., 1986, Slifka et al., 2000), none of those studies reported the death of mice after the treatment. A single study showed T cell depletion by anti-ASGM1 and also addressed the effect of the treatment on the health outcome (Moore et al., 2008). There, anti-ASGM1 treatment was associated with a higher parasite burden after infection with RSV; however, all mice survived and had lower weight loss than control mice, suggesting that ASGM1+ cells also contributed to RSV-induced illness (Moore et al., 2008). Whether this health outcome was caused by depletion of ASGM1+ T cells or NK cells was not specified. The lethality of anti-ASGM1 treatment during secondary *T. gondii* infection suggested that ASGM1+ T cells could be more essential for protection against infection with *T. gondii* parasites than viruses.

ASGM1 could be a marker of specific T cell subsets. In naïve unimmunized mice ASGM1 was expressed on Tcm (defined as CD44^high^, CD62L+, CCR7+) CD8+ T cells (Kosaka et al., 2007). We found that in *T. gondii-*immunized mice ASGM1 was expressed on memory phenotype CD8+ T cells, including both Tem (CD44^high^ CD62L-) and Tcm (CD44^high^ CD62L+) (Srivastava et al., 2018). After reinfection, ASGM1 was distributed between all CD8+ T cell subsets, including Tef and Tn. Importantly, effector phenotype CD8+ T cells (Tef and Tem) were more susceptible to anti-ASGM1 treatment that Tn and Tcm. A previous study also suggested that effector or memory T cells could be more susceptible to the treatment than naïve T cells (Stitz et al., 1986). Virus-specific T cell activity was reduced when anti-ASGM1 was administered after and not before infections with vaccinia or LCMV (Stitz et al., 1986). Furthermore, we found that anti-ASGM1 treatment selectively eliminated CD8+ T cells that provided effector functions (IFNγ and GranzymeB) in response to reinfection. Similar to our findings, the majority of ASGM1+ CD8+ T cells produced IFNγ and were virus-specific after viral infections (Slifka et al., 2000, Moore et al., 2008). It is possible that ASGM1 could be expressed on *T. gondii*-specific CD8+ T cells. In support of this, we found that ASGM1 was enriched on CD8+ T cells that expressed surrogate markers of *T. gondii*-specificity (CD11a and CD49d) (Hwang et al., 2016). Whether or not ASGM1 is a marker *T. gondii*-specific CD8+ T cells would be important to confirm using MHC tetramer staining (Marple et al., 2017). Alternatively, ASGM1 could be expressed on cytokine-activated T cells. ASGM1+ CD8+ T cells from naïve mice were shown to produce more IFNγ than ASGM1-CD8+ T cells after IL-12 (+IL-2) stimulation (Kosaka et al., 2007). After *T. gondii* infection IL-12 is abundantly produced (Gazzinelli et al., 1993, Khan et al., 1994, Hunter et al., 1994, Hunter et al., 1995b). Therefore, IL-12 could induce activation of ASGM1+ CD8+ T cells after *T. gondii* infection. Elucidation of the stimuli necessary for induction of ASGM1 expression on T cells could provide the cues on its functional importance and reveal its potential in use for immunotherapies.

CD4+ T cells were negatively affected by anti-ASGM1 treatment of *T. gondii*-immunized mice. In addition to CD8+ T cells, total splenic and IFNγ-producing peritoneal CD4+ T cells were reduced after the treatment. This is in accordance to an early study on NK cell role in protection against *T. gondii* reinfection in the absence of CD8+ T cells (Denkers et al., 1993). *T. gondii*-immunized CD8+ T cell deficient mice (*β2m-/-)* survived lethal reinfection and died after anti-NK1.1 or anti-ASGM1 treatments. Interestingly, mice died faster after anti-ASGM1 treatment as compared to anti-NK1.1. This result implied that these treatments could have different effects on the cells other than CD8+ T cells and NK cells, potentially CD4+ T cells (Denkers et al., 1993).

The functional role of ASGM1-mediated signaling in T cells is not known. One possibility is that ASGM1 could be functionally relevant for T cell biology as a part of lipid rafts. ASGM1 is a glycolipid that is enriched in lipid rafts, small dynamic ordered domains that compartmentalize cellular processes (Pike, 2006). Upon T cell activation, TCR engagement leads to an aggregation of these domains (Dinic et al., 2015). This aggregation then facilitates TCR signaling by bringing molecules needed for TCR closer to each other. It was found earlier that engagement of GM1 by cholera toxin leads to an activation of T cells (Janes et al., 1999). ASGM1 is a derivative of GM1 and colocalizes with GM1 in the lipid rafts on T cells (Moore et al., 2008). ASGM1 could also be involved in TCR signaling and T cell activation. Whether the presence and quantity of ASGM1 in lipid rafts affects T cell functionality is not known. In addition, whether T cells with enriched ASGM1 lipid raft content have a superior ability for TCR signaling and activation as compared to ASGM1 negative T cells remains to be investigated. That would be important to address as it could be useful for the future design of T cell-based immunotherapies.

As a further point, the presence of ASGM1 could be advantageous for T cell survival during *T. gondii* infection. This is because ASGM1 is an asialylated derivative of GM1 (Moore et al., 2008). Due to an abundance of sialic acids on the cell surfaces, many viruses and bacteria use host-sialylated structures for binding and recognition (Schwegmann-Wessels et al., 2011, Neu et al., 2011). Apicomplexan parasites, including *T. gondii* and *Plasmodium falciparum*, also employ recognition of sialic acids as part of interactions with the hosts (Monteiro et al., 1998, Blumenschein et al., 2007, Gaur and Chitnis, 2011). The *T. gondii* microneme proteins bind to sialylated glycoconjugates on host cells and facilitate invasion (Blumenschein et al., 2007, Friedrich et al., 2010). As any other nucleated cells, T cells can get *T. gondii*-infected and killed by parasites or lymphocytes. Therefore, one possible mechanism by which ASGM1+ CD8+ T cells could have an advantage in survival over ASGM1-CD8+ cells during *T. gondii* infection could be their reduced sialic acid content and thus resistance to being infected by the parasites.

In summary, our data showed that anti-ASGM1 antibody treatment is not a specific approach to study the role of NK cells during *T. gondii* infection. This method could be informative when used with caution and in combination with other NK cell specific methods. We demonstrated that in *T. gondii-*immunized mice certain subsets of CD8+ and CD4+ T cells express ASGM1 and get depleted by anti-ASGM1. Elimination of T cells correlated with increased susceptibility to secondary *T. gondii* infection.

## Materials and Methods

### Mice

C57BL/6 (B6) mice were purchased from The Jackson Laboratory. Animals were housed in specific pathogen-free conditions at the University of Wyoming Animal Facility.

### Ethics Statement

This study was carried out in strict accordance following the recommendations in the Guide for the Care and Use of Laboratory Animals of the National Institutes of Health. The protocol was approved by the Institutional Animal Care and Use Committee (IACUC) of the University of Wyoming (PHS/NIH/OLAW assurance number: A3216-01).

### T. gondii parasites, infections

Tachyzoites (tach.) of RH and RHΔ*cpsII* (*cps1-1,* CPS) strains (kindly provided by Dr. Bzik, Dartmouth College, NH) were cultured by serial passage in human fetal lung fibroblast (MRC5) cell monolayers in complete DMEM (supplemented with 0.2 mM uracil for *cps1-1* strain). The parasites were purified by filtration through a 3.0-µm filter (Merck Millipore Ltd.), then washed with phosphate-buffered saline (1 × PBS) and administered intraperitoneally (i.p.) 1 × 10^3^ RH tach. or 1 × 10^6^ *cps1-1* tach. The brains of 5 wk ME49 infected CBA mice were harvested and 100 ME49 cysts were administered intragastrically (i.g.).

### Real-time PCR for parasite burden

DNA was extracted from entire PECs and spleens harvested from infected mice using a Qiagen DNeasy Blood & Tissue Kit (Qiagen Sciences). Parasite DNA from 600 ng of PECs, 800 ng of splenic or brain tissue DNA was amplified using primers specific for the *T. gondii* B1 gene (forward primer GGAACTGCATCCGTTCATG and reverse primer TCTTTAAAGCGTTCGTGGTC) at 10 pmol of each per reaction (Integrated DNA Technologies) by real-time fluorogenic PCR using SsoAdvanced^TM^ Universal IT SYBR® Green SMx (BIO-RAD) on a CFX Connect^TM^ Real-Time System cycler (BIO-RAD). Parasite equivalents were determined by extrapolation from a standard curve.

### In vivo treatments

Anti-NK1.1 (clone PK136, Bio × cell) was diluted in 1X PBS and administered i.p. at 200 µg one day before (d −1), the day of infection (d 0) and every other day after for the maximum duration of 21 d. Anti-Asialo GM1 (Rabbit, FUJIFILM Wako) was diluted in 1 mL sterile H2O and administered i.p. at 50 µl on d −1, 0 and every third day after.

### Flow cytometry

Peritoneal exudate cells (PEC) and splenocytes were harvested as previously described (Ivanova et al., 2016) and plated at 0.5-1.5 × 10^6^ cells/well. For surface staining, cells were washed twice with 1X PBS and stained for viability in 1 × PBS using Fixable Live/Dead Aqua dye (Invitrogen) for 30 min. After washing with 1 × PBS, surface staining was performed using antibodies diluted in stain wash buffer (SWB, 2% FBS in 1 × PBS and EDTA) for 25 min on ice. For functional assays, cells were stimulated with 20 µg/mL Toxoplasma Lysate Antigen (TLA) for 8 h in complete Isclove’s media and then treated with 0.7 × PTIC for 4 h. For additional functional assays, cells were incubated for 8 h in complete Isclove’s media and then treated with 0.7 × Cell Stimulation Cocktail containing PMA/ionomycin and PTIC (eBioscience, ThermoFisher) for 4 h. After live dead and surface staining, cells were fixed and permeabilized for 1 h on ice (BD bioscience, Fix/Perm solution) followed by intracellular staining in 1 × permeabilization wash buffer (BD) with anti-IFNγ, anti-TNFα, anti-Granzyme B antibody for 40 min. After washing twice with 1X PBS, cells were resuspended in 1X PBS and acquired using Guava easyCyte HT (Millipore). All samples were analyzed with FlowJo software (Tree Star). The antibodies purchased from Biolegend were: CD3 (17A2), CD4 (RM4-5), CD8b (YTS156.7.7), NK1.1 (PK136), Asialo-GM1 (Poly21460), CD44 (IM7), CD62L (MEL-14), CD11a (M17/4), CD49d (R1-2). The antibodies purchased from eBioscience/ThermoFisher were: IFNγ (XMG1.2), TNFα (Mab11), Granzyme B (NGZB).

### Statistical analysis

Statistical analysis was performed using Prism 7.0d (GraphPad) or Microsoft Excel 2011. Significant differences were calculated using either unpaired Student’s t-test or analysis of variance (ANOVA). The log-rank (Mantel-Cox) test was used to evaluate survival rate. Significance is denoted where *p<0.05, **p<0.01, ***p<0.001, ****p<0.0001.

## Acknowledgments

We thank Rida Fatima, Tiffany Mundhenke, Sally Murray and Ryan Krempels for assistance.

## Notes

This work is supported by grants from the American Heart Association AHA 17GRNT33700199 and University of Wyoming INBRE P20 GM103432 DRPP awarded to JPG. DLI is a University of Wyoming INBRE P20 GM103432 Graduate Assistantship recipient. SLD is a University of Wyoming INBRE P20 GM103432 supported undergraduate fellow. This project is supported in part by a grant from the National Institute of General Medical Sciences (2P20GM103432) from the National Institutes of Health. The content is solely the responsibility of the authors and does not necessarily represent the official views of the National Institutes of Health

